# Nanoscopy of Organelle Handoff Portals Reveals Direct Coupling between ER Remodeling and Microtubule-Based Transport

**DOI:** 10.1101/2025.09.18.677214

**Authors:** Jin-Sung Park, Il-Buem Lee, Hyeon-Min Moon, Hyeonjun Jeon, MinHyeong Lee, Chungho Kim, Seok-Cheol Hong, Minhaeng Cho

## Abstract

How biosynthetic organelles leave the endoplasmic reticulum (ER) and engage with microtubule tracks remains a central question. Combining interferometric scattering with fluorescence nanoscopy, we tracked nanometer-scale handoff events in living cells. ER-derived organelles undergo biased diffusion along ER tubules toward nearby microtubules. ER three-way junctions function as nanoscopic hubs where cargo pauses, contacts multiple microtubules, and then launches onto a track for long-range travel. Remarkably, the ER maintains a membrane tether to the departing cargo, extending its tubules and forming new junctions, thereby coupling inter-network transfer with membrane morphogenesis. These observations reveal an integrated mechanism that links organelle biogenesis, directional trafficking, and continual ER- and cellular remodeling, underscoring the ER’s active role in steering transport and repurposing its own output within the crowded intracellular environment.

## INTRODUCTION

The endoplasmic reticulum (ER) is a central hub in the cellular transport system, responsible for synthesizing, folding, and trafficking proteins and lipids.^1,2^ Newly synthesized proteins are packaged into membrane-bound carriers for subsequent processing and sorting.^3^ The ER also contributes to *de novo* organelle biosynthesis, giving rise to structures such as omegasomes – precursors of autophagosomes (APGs) – as well as peroxisomes and lipid droplets.^4,5^ These organelles are then transported along microtubules, which serve as directional tracks for motor proteins for efficient intracellular trafficking. Extensive studies have examined ER interactions with organelles and the cytoskeleton, highlighting their roles in maintaining cellular organization and function.^6-13^ Despite their distinct physical and functional characteristics, the ER and microtubules work in coordination to regulate organelle positioning, transport, and signaling.^6,9,14^ Although this interplay is presumed to be vital for the transport of ER-associated organelles, the underlying mechanisms – such as how organelles translocate between the ER and microtubules or navigate toward nearby microtubules for long-range transport – remain poorly understood.

Here, we present a direct visualization of organelle transfer from the ER to microtubules, occurring within their spatially overlapping networks. To capture these nanoscopic cellular events, we employed a custom-tailored imaging platform based on interferometric scattering (iSCAT) microscopy integrated with fluorescence microscopy (Figure S1). Using only a single-color label, this system enabled simultaneous visualization of cytoplasmic ER and microtubule networks, along with target organelles, at high spatiotemporal resolution. Our observations show that the ER plays an active role in biasing the directional diffusion of organelles toward neighboring microtubules. During this process, ER three-way junctions act as transfer points, transiently anchoring organelles and forming contact sites with multiple microtubules until handoff occurs. We further find that the ER actively engages in its own remodeling by keeping its tips tethered to organelles when the organelles are transferred to microtubules and subsequently transported along the microtubule networks, which may constitute a fundamental principle that inevitably brings the two networks into spatial congruence and provides a mechanistic basis for their intimate functional coordination.

## RESULT AND DISCUSSION

### ER Remodeling Induced by Direct Collision of ER Tubules by a Directionally Moving Cargo

The ER forms an interconnected network of perinuclear membrane sheets and tubules. Although it is most readily visualized in the quasi-2D periphery of cultured cells, it can also be observed alongside other cytoplasmic organelles, such as mitochondria and lipid droplets, using phase-enhanced optical microscopy techniques.^15-17^ iSCAT microscopy has since expanded its applications across diverse research fields, including nanoparticle detection^*18-21*^ and cell biology.^22-27^ Recently, iSCAT-based approaches have enabled high-contrast visualization of the nanometric topography of the tubular ER network by incorporating optical modules such as confocal pinholes and rotating diffusers to suppress speckle-like background noise originating from the complex 3D cellular environment.^*28-30*^

Notably, we successfully captured these tubular ER structures in a single iSCAT snapshot even without spatial filtering (Figure S2A), although additional image processing is typically required to enhance their contrast (Experimental Section and Figures S2B–D). In static background removed (SBR-)iSCAT images, the contrast is improved by suppressing speckle background noise, but ER structures often remain obscured due to interference from other cytoplasmic components (Figures S2C and 1A). Given the highly dynamic nature of the ER,^9^ we employed time-differential (TD-)iSCAT imaging to suppress relatively static cytoplasmic signals and highlight dynamic ER remodeling and associated cargo movements (Figures S2D and 1B).^26,27^ This dynamic network is further resolved using an iSCAT variance map, generated by calculating the pixel-wise standard deviation of TD-iSCAT images over time (Experimental Section and Figure 1C), in the light of a related approach reported previously.^25^ The resulting variance map, which is particularly sensitive to dynamic structures such as the ER and mobile cargo, closely correlates with the complementary fluorescence image of labeled ER (Figure 1D and Movie S1). In the sections that follow, fluorescent LC3 was used instead to selectively visualize APGs and microtubule filaments.

**Figure 1.**
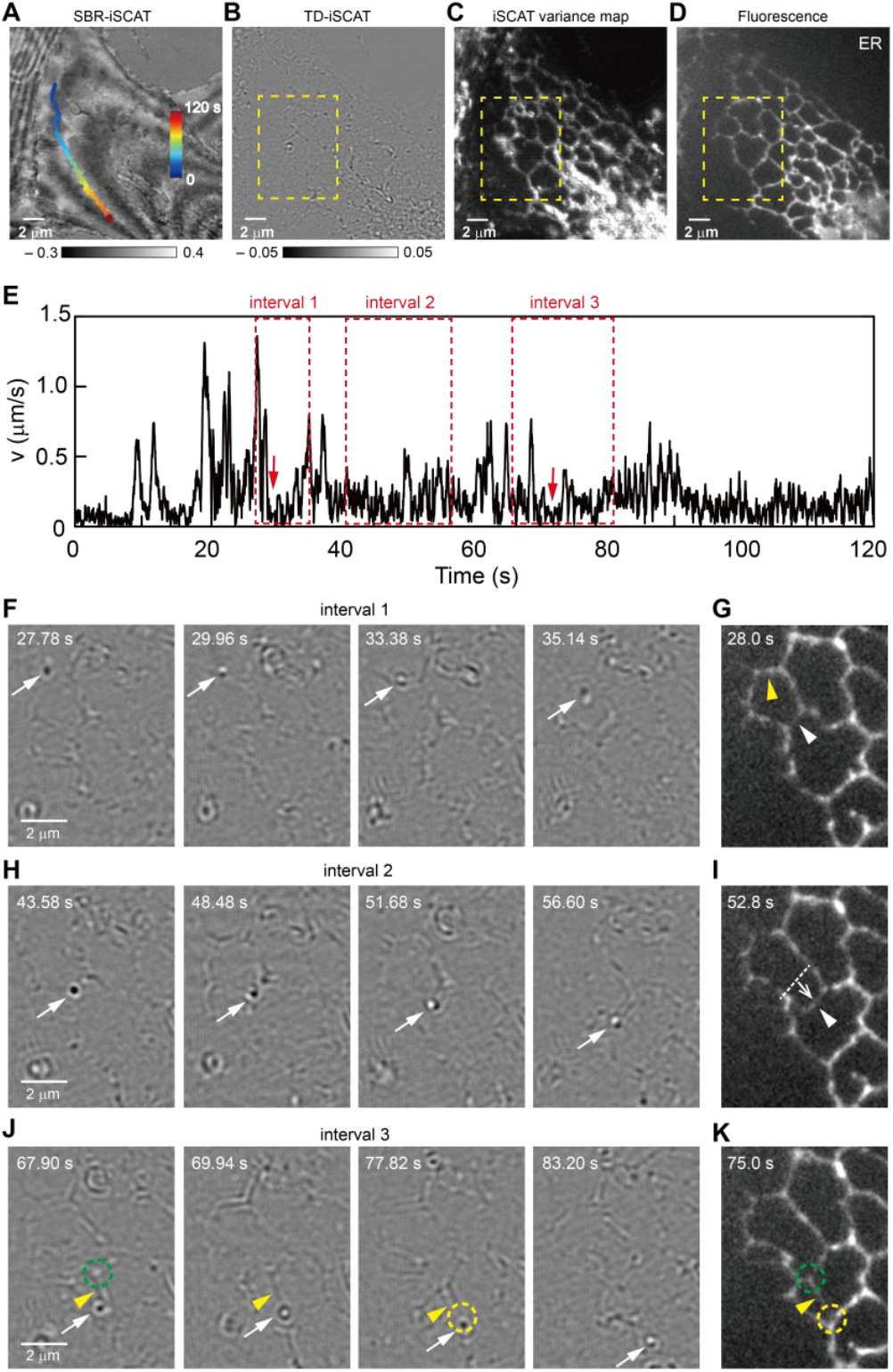
Label-free visualization of ER-cargo interaction. (**A**) SBR-iSCAT image revealing various subcellular structures in a GFP-ER labeled COS-7 cell with one representative cargo trajectory overlaid and color-coded by time. (**B**) TD-iSCAT image highlighting dynamic ER networks and mobile cargos. In (A and B), the grayscale legends represent iSCAT contrasts. (**C**) iSCAT variance map generated by calculating the standard deviation of TD-iSCAT in each pixel over a 200-frame TD-iSCAT sequence (4 s). (**D**) Complementary fluorescent image showing the ER networks. See Movie S1. (**E**) The υ of the tracked cargo shown in (A), smoothed with a 50-point (1 s) moving average. (**F, H, and J**) TD-iSCAT image sequences showing three notable events of ER-cargo interactions (F: collision of a moving cargo with a transverse ER tubule; H: collision of the cargo with another transverse ER tubule and its displacement thereby; J: formation of an ER tether by the cargo (hence, generation of an ER three-way junction, as indicated by green circle in the 1^st^ image) and fusion of the tether to a third transverse ER tubule (hence, generation of another ER three-way junction, as indicated by yellow circle in the 3^rd^ image) from the yellow-boxed area in (B). (**G, I, and K**) Complementary fluorescence images showing the ER structures involved in the ER-cargo interaction events shown in (F, H, and J). In (K), the green and yellow circles mark the same sites indicated in (J). For (F, H, and J), see Movies S2 to S4.

The results in Figure 1 demonstrate that iSCAT provides valuable insights into the interaction between the ER and dynamic cargos. The instantaneous speed (*v*) of a cargo, tracked from the SBR-iSCAT image sequence acquired at 50 Hz and overlaid in Figure 1A, exhibits pronounced fluctuations (Figure 1E). Our analysis reveals that direct ER-cargo interactions constitute a major source of these variations. In interval ‘1’ (Figures 1E,F, and Movie S2), the cargo (white arrow in Figure 1F) undergoes a head-on collision with a transversely oriented ER tubule at *t* = 29.96 s, which is part of a polygonal ER network (yellow arrowhead in Figure 1G), leading to a sharp drop in *v* (red arrow in interval ‘1’ marked in Figure 1E). The speed recovers after the cargo crosses the ER tubule at *t* = 35.14s. Such collisions not only slow the cargo but also induce ER remodeling. In interval ‘2’ (Figures 1E,H, and Movie S3), at *t* = 43.58 s, the cargo pushes against another ER tubule, displacing it in the direction of motion (white arrowheads in Figures 1G,I). Subsequently, in interval ‘3’ (Figures 1E,J, and Movie S4), a new ER tubule extends along the cargo’s path (yellow arrowheads in Figures 1J,K) after forming a new ER three-way junction (green circle in Figures 1J,K). This new tubule then connects to a downstream ER tubule, generating a second ER three-way junction (yellow circle in Figures 1J,K).

These observations support the concept of ‘ER hitchhiking’, in which the ER dynamically remodels itself by harnessing cargo movement along the cytoskeleton – an essential mechanism for ER network reorganization.^*9,26,31-34*^ Moreover, they provide a physical mechanism that naturally produces—and accounts for—the highly correlated alignment of ER tubules with microtubules and the frequent spatial overlap between ER three-way junctions and microtubules documented in earlier studies.

### APG Handoff Sequence Visualized across ER Three-Way Junction–Microtubule Crossroads

To investigate how ER-associated organelles translocate to microtubules for long-range transport, we used COS-7 cells expressing GFP-labeled LC3, a microtubule-associated protein light chain that serves as a marker for APGs, our ER-derived model cargos (Experimental Section).^35^ The ER initiates APG biogenesis by promoting the nucleation of the phagophore, which subsequently expands to engulf autophagic cargos and ultimately seals to form a mature APG.^36^ Consistent with this process, we observed that LC3 fluorescence typically appeared as puncta (Figure S3A), emerging from an elongated membrane segment (white-boxed area at *t* = 0 s) and transforming into a punctum after 16 s (inset in Figure S3A and Movie S5). Although LC3-labeled APGs were also visible in both SBR- and TD-iSCAT images (Figures S3B,C), they could not be reliably distinguished from other cargos of similar morphology without complementary fluorescence imaging. Intriguingly, we observed ER hitchhiking behavior, in which a newly extended ER tubule elongates by tethering to an APG undergoing directional transport, thereby driving ER remodeling (Figure S3D).

LC3 is also distributed along microtubules, making it a useful marker for visualizing both microtubules and APGs within cells.^37^ In fluorescence images, APGs typically appear as dot-like or spherical structures, while microtubules are seen as filamentous networks (Figure 2A). By acquiring images simultaneously with iSCAT and fluorescence microscopies, we could examine APG transports in relation to both the ER and microtubule cytoplasmic networks (Figures 2A–C and Movie S6). While fluorescence imaging selectively highlights APGs (inset in Figure 2A, corresponding to the white-dashed area in Figure 2A), iSCAT and its variation map capture both APGs and other native cargos (insets in Figures 2B,C, and upper inset in Figure 2D, corresponding to the white-dashed area in Figures 2B–D, respectively). The appearance of these cargos, however, differs between SBR- and TD-iSCAT images due to differences in image processing (insets in Figures 2B,C): SBR-iSCAT emphasizes static subcellular structures, whereas TD-iSCAT highlights only dynamic features with greater detection sensitivity (Experimental Section).^26^ In the iSCAT variance map (Figure 2D), ER three-way junctions – highlighted by arrowheads at the vertices of the polygonal ER network (bottom inset in Figure 2D, corresponding to the yellow-dashed area) – serve as direct contact points and potential stalling sites for numerous cargos, including APGs. Among at least six cargos identified in the bottom inset of Figure 2D, only one of them – indicated by the red arrowhead – was confirmed as an APG, as shown by the red-dashed circle in Figures 2A and C, based on fluorescence from the corresponding area.

**Figure 2.**
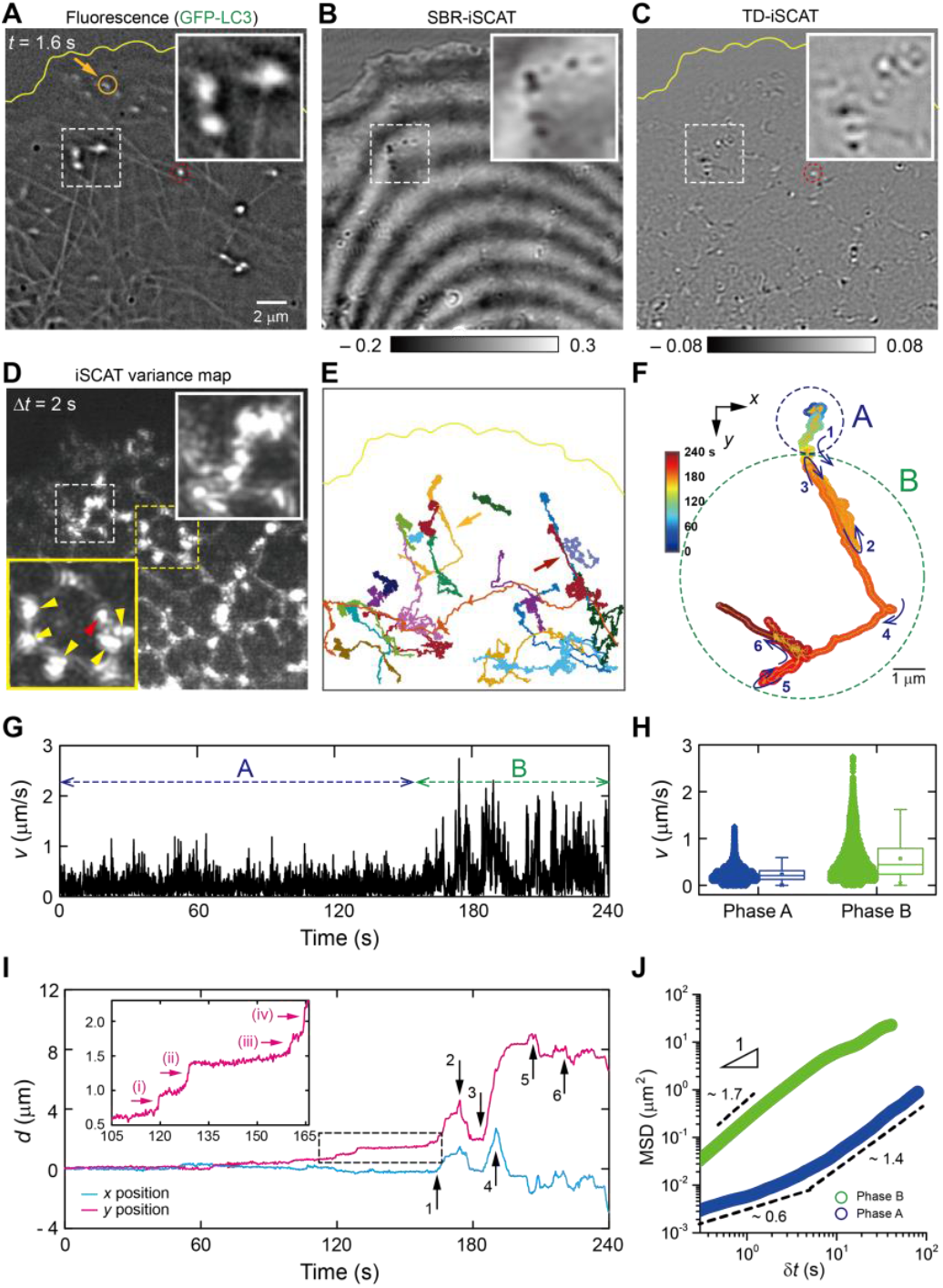
Heterogeneous APG movements over highly intricated cellular networks. (**A**) Fluorescence image showing LC3-labeled APGs and a complex network of microtubules therewith. (**B**) SBR-iSCAT snapshot revealing cytoplasmic structures, including APGs. (**C**) TD-iSCAT snapshot highlighting dynamic cargos and ER networks. In (B and C), the grayscale legends represent iSCAT contrasts. (**D**) iSCAT variance map showing the spatial distribution of ER networks and associated cargos. Insets in (A to C) and the upper inset in (D) display the zoomed-in views of the white-dashed area showing the locations of APGs and other unlabeled cargos. The bottom inset in (D) displays a zoomed-in view of the yellow-dashed area showing cargos (arrowheads) sticking around ER three-way junctions. Among them, only one cargo is specified as an APG by fluorescence (red circles in (A and C)). (**E**) Trajectories of 21 APGs extracted from a fluorescence image sequence acquired at 10 Hz. (**F**) One representative APG trajectory, indicated by an orange arrow in (A and E), extracted from the corresponding SBR-iSCAT image sequence acquired at 200 Hz with arrows indicating directional changes and turning points. (**G**) Variation in υ calculated from the trajectory in (F) and smoothed by 100-point moving average (*Δt* = 0.5 s). (**H**) Boxplot of υ values for two distinct phases, ‘A’ and ‘B’, in (F and G). (**I**) Variations in *x*- (blue) and *y*-displacements (red) from the trajectory in (F), with arrows marking the turning points. Inset: zoomed-in view of the dashed-boxed region showing three short segments of directionally drifting motion, (i-iv). (**J**) MSD *vs*. time plots for two distinct modes of APG motion (blue: phase ‘A’ and green: phase ‘B’). For (A to C), see Movie S6.

We tracked the motion of 21 individual APGs using time-lapse fluorescence imaging at 10 Hz (Figure 2E). Individual trajectories for all 21 APGs, along with corresponding fluorescence snapshots, and simultaneously acquired iSCAT images are shown in Figure S4. To conduct a detailed investigation of the trajectory of a representative APG (trajectory 1 in Figure S4), indicated by an orange arrow in Figures 2A,E, we retrieved its trajectory from the corresponding SBR-iSCAT image sequence, which was acquired at a higher frame rate of 200 Hz (Figure 2F). This trajectory reveals two distinct dynamic phases, labeled ‘A’ and ‘B’, characterized and distinguished by their velocity (*v*) and variability (Figures 2G,H): phase ‘A’ is characterized by low-speed one-dimensional diffusion, whereas phase ‘B’ exhibits (bi-)directional motion with abrupt directional reversals and turns at elevated speeds. Abrupt directional changes, including U-turns, are marked by numbered arrows (1 ∼ 6), which correspond to sharp shifts in displacement along the *x* and *y* directions (Figures 2F,I, and 3A,D). Intriguingly, phase ‘A’ shows one-dimensional anomalous diffusion, transitioning at *δt* ∼ 5 s from sub-diffusive to super-diffusive behavior as the APG gradually drifts in a preferred direction (blue curve in Figure 2J). By contrast, phase ‘B’ exhibits directed runs punctuated by intermittent to-and-fro reversals, yielding a higher (α ≈ 1.7) diffusion exponent (green curve in Figure 2J), although the MSD curve becomes less steep beyond *δt* ∼ 10 s, reflecting occasional divergence of the APG at microtubule intersections. These dynamic changes suggest that the APG undergoes translocation from the ER to a microtubule, prompting a closer investigation of the translocation process.

### ER-Guided Translocation of APGs to Nearby Microtubules

Next, we examined in detail how the APG translocates from the ER to a microtubule by analyzing its trajectory in Figure 2F, in relation to the underlying ER and microtubule networks. In region ‘1’ of the trajectory (Figure 3A), the APG – marked by yellow-dashed circles in both the SBR- and TD-iSCAT images (Figure 3B) – appears initially anchored at the tip of an ER tubule branching from a three-way ER junction, which forms part of a polygonal ER structure, as indicated by the purple arrow in the iSCAT variance map (Figure 3B). Microtubules are observed to exist coincidently with this polygonal ER structure presumably due to microtubule-associated ER remodeling: one is aligned along the ER structure, while another crosses it, as marked by the orange arrows and arrowheads, respectively, in the fluorescence image (Figure 3B). These microtubules are overlaid as green solid and dashed curves in the zoomed-in trajectory (Figure 3C).

**Figure 3.**
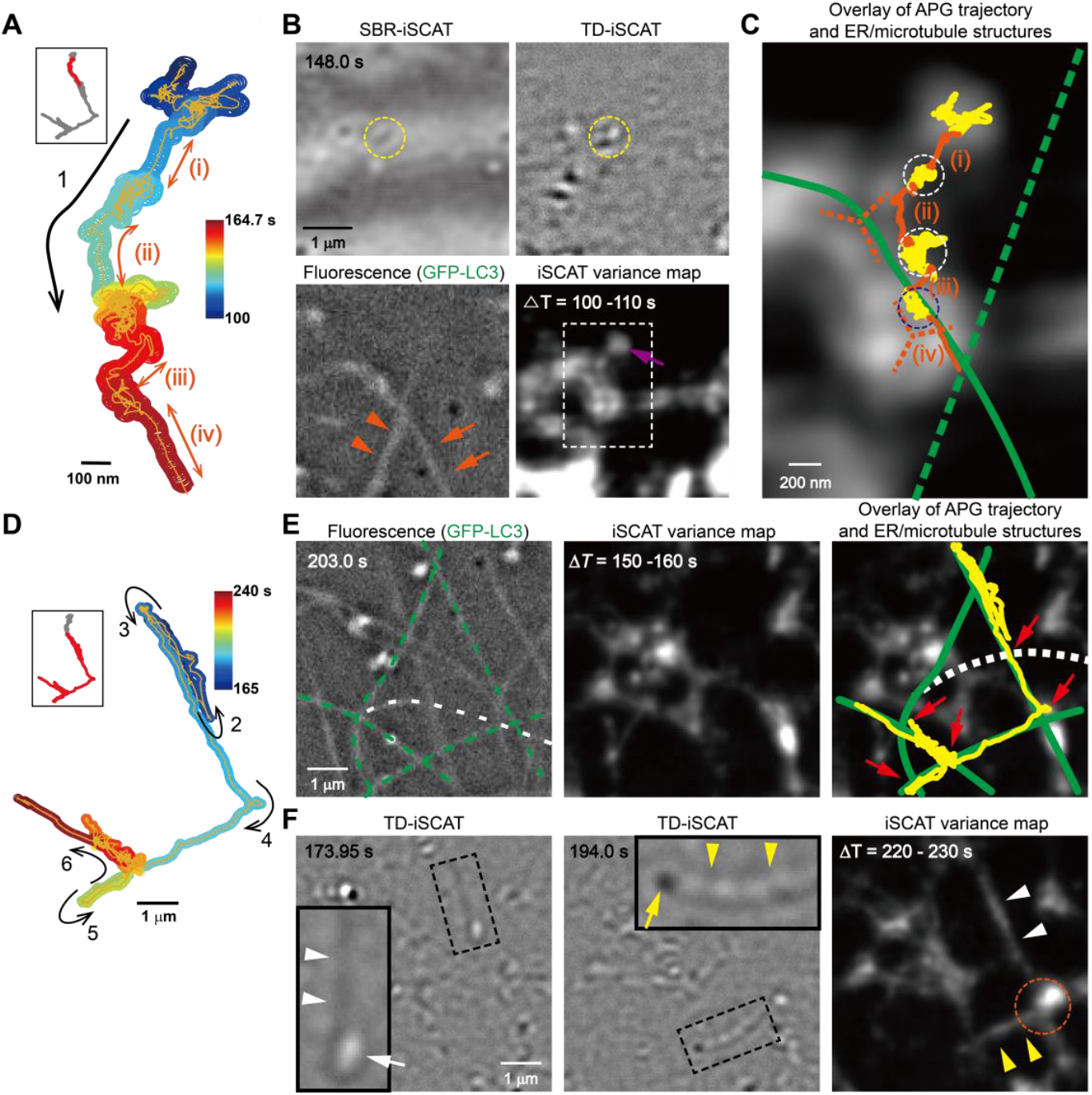
Motion switching of APG from ER-guided migration to microtubule-based transport and ER remodeling thereby. (**A**) Zoomed-in APG trajectory showing ER-guided movement (arrow ‘1’) with directionally drifting regions (i – iv), corresponding to the red curved trace highlighted in the full trajectory (inset). (**B**) SBR-iSCAT, TD-iSCAT, fluorescence snapshots and iSCAT variance map revealing the location of the tracked APG and the underlying microtubule and ER structures. (**C**) Zoomed-in image of the white-boxed region in (B) (microtubules: green solid and dashed curves, indicated by orange arrows and arrowheads in the fluorescence image of (B), respectively; ER three-way junctions: orange-dashed branched symbols). (**D**) Zoomed-in APG trajectory showing the directional movement along microtubules, corresponding to the red branched traces highlighted in the full trajectory (inset). (**E**) Fluorescence image (left), iSCAT variance map (center), and annotated and superimposed iSCAT variance map (right), showing APG passages (yellow) and underlying networks. (**F**) TD-iSCAT snapshots taken at two different time points showing ER tethering to the APG. iSCAT variance map shows newly formed ER tubules (white and yellow arrowheads) and an ER junction (orange circle). Insets show the detailed views of ER-tethered APGs corresponding to the black-dashed boxes in (F).

Comparison of the APG trajectory with the underlying ER and microtubule architecture reveals that the APG moves directionally along the curved edge of the polygonal ER structure (arrow in Figure 3A). This motion is punctuated by intermittent pauses – marked by two white and one blue circles along the trajectory in Figure 3C – interspersed between segments of directed motion, indicated by orange arrows (i)-(iv) in Figure 3A. These correspond to the orange-colored routes (i)-(iv) in the trajectory shown in Figure 3C (see also the inset of Figure 2I). The white-circled pauses appear to be associated with ER three-way junctions, highlighted by orange-dashed branches for visual guidance (Figure 3C).

In contrast, the blue-circled pause likely corresponds to the transfer event in which the APG transitions from the ER to the microtubule. At ER three-way junctions, membrane tension is distributed evenly across the three tubules, creating directional frustration that transiently traps ER-bound cargo. This prolonged dwell time provides an opportunity for efficient capture by microtubule filaments, which are frequently juxtaposed to such junctions owing to their conserved spatial association. Consistent with this interpretation, the APG subsequently moves directionally along the microtubule network, as shown by its trajectory in Figure 2F and the magnified view in Figure 3D.

Among the 21 APGs we tracked in regions where polygonal ER and microtubules coexist (Figures 2E and S4), ER-guided diffusion culminating in an ER-to-microtubule handoff was the predominant behavior (trajectories 1, 5–8, 10–11, 13–14, 16–17). Less frequently, we observed microtubule-to-ER reversals (trajectories 20–21), episodes of confinement within microtubule-encircled ER polygons (trajectories 2–4, 9, 12, 18–19), and, rarely, a long uninterrupted microtubule run (trajectory 15).

Following its handoff to microtubules, the APG exhibits directional movement punctuated by abrupt turns. Compared with its average speed, 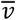, during ER-guided directional drift along polygonal ER edges 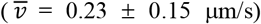, the APG moves significantly faster along microtubules 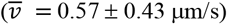 and maintains greater directional persistence (Figures 2H,J). In the trajectory overlaid on the microtubule network, most abrupt turns occur at microtubule intersections (red arrows), as shown in the annotated and superimposed iSCAT variance map (Figure 3E). This suggests that intersections may serve as physical barriers that interrupt the APG’s persistent motion^38,39^ presumably via collision with obstructing microtubule filaments and also strategic checkpoints that diversify the APG’s destinations. Notably, we observed that the ER tip tethered to the APG elongates as the APG travels along the microtubule (insets in the 1^st^ and 2^nd^ images of Figure 3F). This ER tethering promotes the extension of new ER tubules along the microtubule path and facilitates the formation of a new ER junction at the APG’s turning point near the microtubule intersection (orange-circled region in the 3^rd^ image of Figure 3F).

Beyond the *de novo* three-way junctions generated when a cargo laterally collides with an ER tubule (Figure 1), these two additional modalities appear to operate in ER remodeling: (i) tether elongation, in which the bounded cargo pulls and extends its parent tubule, and (ii) junction fusion, whereby the same cargo collides with a distal tubule, bridging the two membranes into proximity and nucleating a new three-way junction. Collectively, these three cargo-driven actions constitute a minimal mechanistic toolkit for ER morphogenesis. Iterative cycles of tether extension, junction budding, and cargo-mediated fusion can transform a simple network of tubules into a reticulate lattice, adjust network connectivity in response to local traffic demands, and couple membrane growth to outbound cargo flow. We propose that the frequency and sequence of these elementary mechanistic modules set the tempo of ER remodeling in living cells.

### Capturing the Moment of APG Transfer at ER-Microtubule Junction

To gain deeper insight into the ER-microtubule interactions during APG transport, we analyzed another APG trajectory (brown arrow in Figure 2E and trajectory 8 in Figure S4) within the context of the underlying microtubule network in a cell body (Figure 4A). Unlike the relatively static APG associated with the ER in a thin lamellipodium (Figure 3A), this APG exhibited more dynamic behavior along the ER network, as evident in the TD-iSCAT image with the overlaid trajectory (Figure 4B) and its corresponding variance map (Figure 4C).

**Figure 4.**
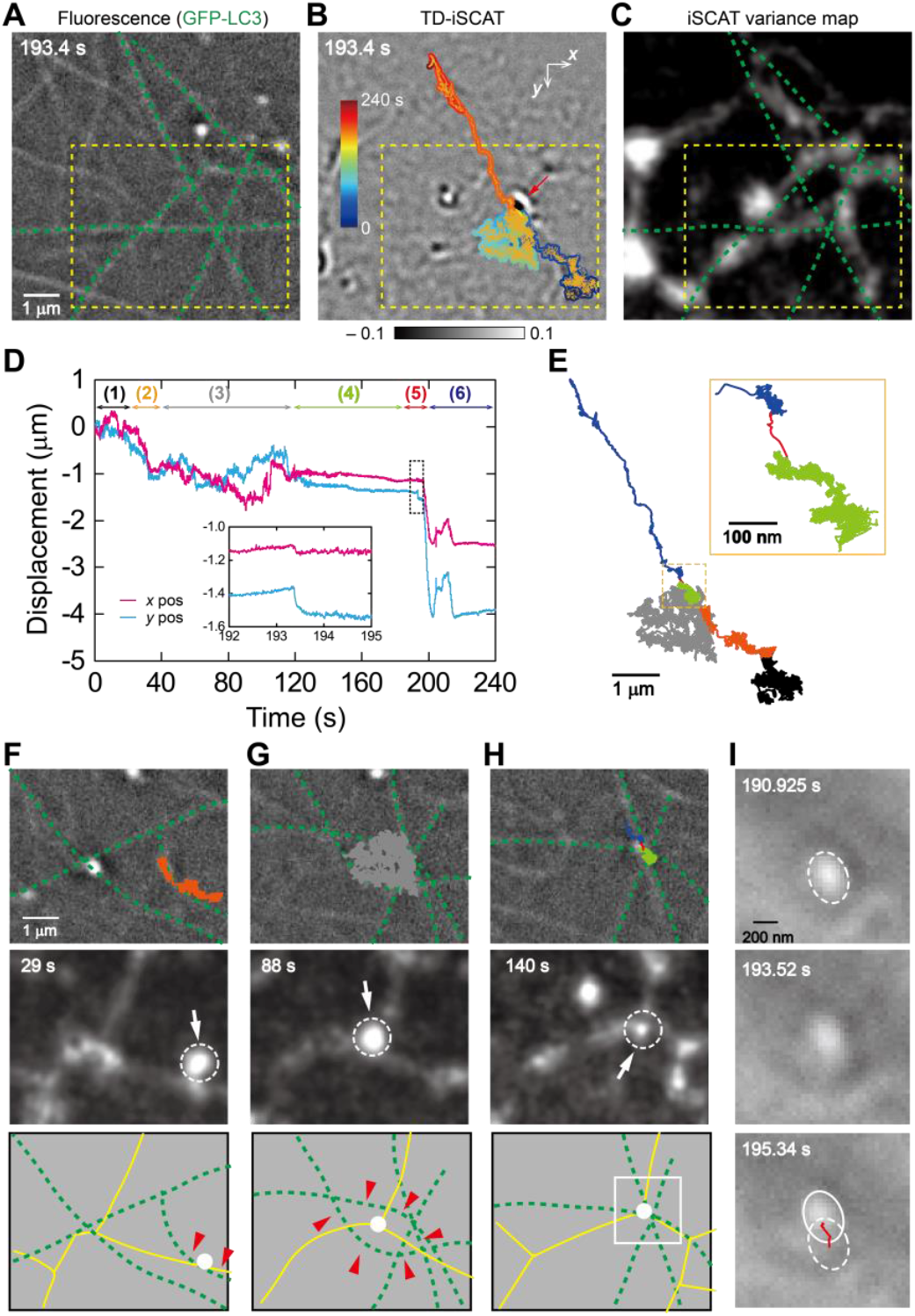
Visualizing the moment of APG transfer at the ER-microtubule junction. (**A**) Fluorescence image showing microtubules and APG locations. (**B**) TD-iSCAT image overlaid with the APG trajectory indicated by the brown arrow in Figure 3D. In (B), the grayscale legends represent iSCAT contrasts. (**C**) iSCAT variance map. (**D**) *x*- and *y-*axis displacement plots derived from the trajectory in (B). Inset highlights the moment of APG transfer from the ER to a microtubule. See Movies S7, S8, and S9, corresponding to the phases of ‘1 - 3’, ‘4 - 5’, and ‘6’. (**E**) Segmentation of the APG trajectory into the 6 different phases marked in (D). The segments are color-coded as labeled in (D). Inset: zoomed-in view of the yellow-boxed area in the trajectory. (**F, G, and H**), Fluorescence images with microtubules highlighted in green (top), iSCAT variance maps (middle), and schematic representations of the ER (yellow) and microtubule (green) networks (bottom) in phases ‘2’, ‘3’ and ‘4’ (yellow dashed boxes in Figures 4A,C). For the dynamic process of ER remodeling during APG transport, see Figure S6 and Movie S11. (**I**) SBR-iSCAT image sequence capturing the moment of APG transfer from the ER to a microtubule. The trajectory segment (red) is overlaid in the bottom image, with outlines indicating the APG position at *t* = 190.925 (dashed circle) and 195.34 s (solid circle). See Movie S10.

Based on the dynamic features of APG motion observed in the *x-* and *y-*displacement plots (Figure 4D), we segmented the trajectory into six distinct phases, color-coded in Figures 4D and E: (1 and 3) random diffusion (black and gray), (2) directional drift (orange), (4) stalled motion with very slow drift (green), (5) instantaneous short-range transport (red), and (6) long-range directional transport (blue). Corresponding time-lapse sequences are provided in Movies S7 (phases 1–3), S8 (phases 4–5), and S9 (phase 6). Initially, the APG underwent random diffusion along the polygonal ER network in a region that also contained microtubules (phase ‘1’ in Figure 4D and black segment in Figure 4E), then transitioned to directional drift (phase ‘2’ in Figure 4D and orange segment in Figure 4E) along an ER segment partially overlapping a curved microtubule (red arrowheads in the bottom panel of Figure 4F). The APG was not transported further along this microtubule; instead, it reverted to random diffusion (phase ‘3’ in Figure 4D and gray segment in Figure 4E) while remaining colocalized with an ER three-way junction and apparently confined by the surrounding microtubule array (Figure 4G and its zoomed-in trajectory color-coded by time in Figure S5). This confined diffusion reflects an unbound or weakly interacting state with adjacent microtubules, while the APG remained associated with the ER. Notably, when the APG-associated ER junction colocalized with multiple microtubules, the APG abruptly stalled and then began to drift slowly toward the ER-microtubule intersection (phase ‘4’ in Figure 4D and green segment in Figure 4E). At this intersection, the APG exhibited a brief burst of directed movement (phase ‘5’ in Figure 4D and its inset, red curved route in the inset of Figure 4E, and Movie S10) - marking the moment of transfer at the ER-microtubule junction. Concomitantly, we observed cargo-driven ER remodeling around the APG, indicating dynamic interplay between the cargo and the surrounding ER network (Figure S6 and Movie S11).

These dynamic characteristics of the APG trajectory are also captured in the MSD curves (Figure 5). In phases ‘1’ and ‘3’ (black and gray segments in Figure 4E), the diffusion exponents (α) are close to 1, indicating normal Brownian motion. In phase ‘2’ (orange), α likewise remains near 1, not because directional motion is absent, but because the short duration of the event (∼ 7 s, from 25 to 32 s) restricts the MSD time window, making the underlying directional drift less evident. In contrast, phase ‘4’ (green) shows a crossover in the the MSD curve from sub-diffusion (α ≈ 0.2) to super-diffusion (α ≈ 1.6) at *Δδt* ≈ 4 s, suggesting a transition from restricted diffusion to slow drift at a stably formed ER-microtubule intersection, where the APG becomes temporarily stalled. As expected, in phase ‘6’ (blue), where the APG was transported along microtubules, α approaches 2.

**Figure 5.**
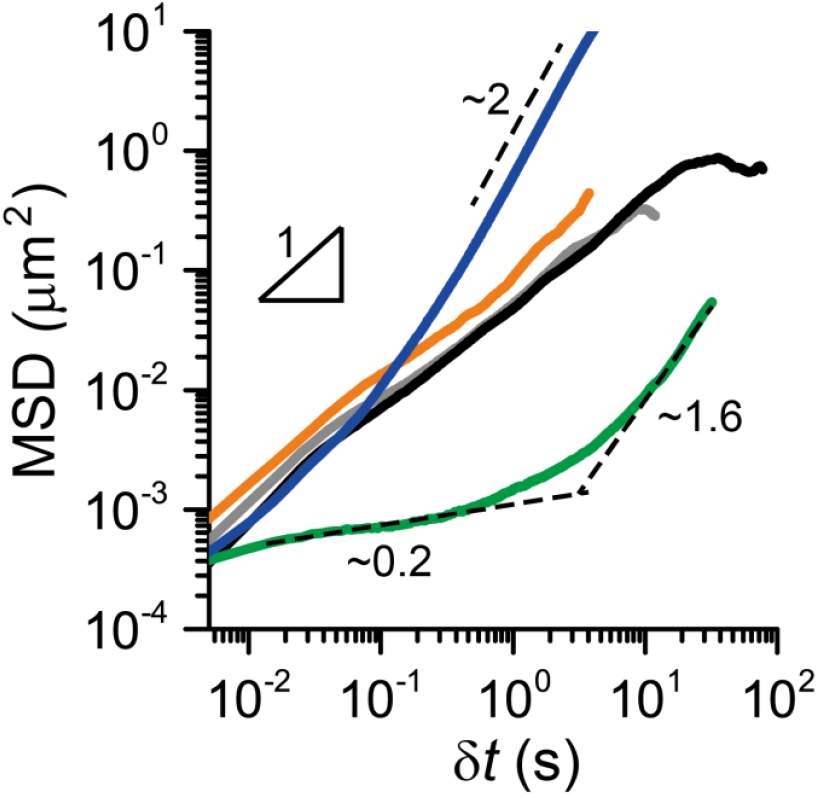
MSD analysis revealing dynamic characteristics in six distinct regions of the APG trajectory. In phases ‘1’, ‘2’ and ‘3’ (black, orange, and gray) in Figure 4E, the APG exhibits normal diffusion with α ≈ 1. In contrast, phase ‘4’ (green) shows anomalous diffusion, characterized by a crossover from sub-diffusion (α ≈ 0.2) to super-diffusion (α ≈ 1.6) near the time interval of *δt* ≈ 4 s. Region ‘6’ (blue) displays a typical directed movement with α ≈ 2.

## CONCLUSION

By simultaneously visualizing organelle transport alongside the underlying ER and microtubule networks, we directly captured organelle transfer events at ER-microtubule intersections. Our observations reveal that the ER steers the directional, mesh-confined diffusion of ER-associated organelles, such as APGs, toward nearby microtubules, thereby facilitating their transfer. ER three-way junctions serve as key stopover sites in this translocation process, stably anchoring organelles while forming physical contacts with multiple microtubules.

The dynamic behavior of APGs alternates between normal and anomalous diffusion, depending on their interaction with the ER and microtubules. At ER-microtubule intersections, APGs transition from sub-diffusive to super-diffusive motion upon the formation of dynamically stable junctions. While the spatial overlap between the ER and microtubules has been reported previously,^40,41^ the mechanism orchestrating this overlap has remained unclear. Our findings show that the ER remodels itself by tethering the tip of an ER tubule to ER-synthesized organelles as they travel along microtubules, thereby promoting the formation of overlapping ER and microtubule networks. Furthermore, ER tubules extended through organelle hitchhiking can give rise to new ER three-way junctions at microtubule intersections, further reinforcing this spatial overlap. Collectively, these results highlight the critical role of ER-microtubule interplay in facilitating efficient organelle transport and driving dynamic ER remodeling.

## EXPERIMENTAL SECTION

### Cell Culture Procedures

COS-7 cells were cultured in a 35-mm confocal dish (Cat. No. 101350, SPL, Korea) pre-coated with 0.1 mg/ml poly-D-lysine (P7886, Sigma-Aldrich, USA). An optimal (low) seeding density of approximately 3 × 10^4^ cells per dish was used for iSCAT imaging of isolated single cells. Cells were incubated at 37 °C in a humidified 5% CO_2_ and 95% air atmosphere in Dulbecco’s modified eagle medium (DMEM) (Gibco, USA) supplemented with 10% fetal bovine serum (FBS, Gibco, USA) and 1% penicillin/streptomycin (Gibco, USA). To evaluate the capability of iSCAT for label-free detection of ER dynamics (Figure 1), COS-7 cells expressing GFP-labeled ER were prepared by adding 6 μL of BacMam 2.0 reagent (Cat. No. C10590, Invitrogen, USA) 24 hours after cell seeding, followed by incubation for 16 hours.

For APG visualization experiments, COS-7 cells were transduced with a lentivirus encoding GFP-LC3 for fluorescent labeling of APGs, as previously described.^31^ Transduction efficiency was confirmed to be 100% based on GFP fluorescence. COS-7 cells expressing GFP-labeled LC3 were imaged with fluorescence-integrated iSCAT microscopy for dual-mode detection. To control the temperature and pH condition of the sample dish for long-term iSCAT live-cell imaging, the sample dish was placed in a mini-incubating chamber (Chamlide, Live Cell Instrument, Korea) mounted on a piezo stage (MZS500, Thorlabs, USA).

### Experimental Setup of iSCAT Microscopy

The sample was illuminated by raster scanning (Figure S1) with a continuous-wave diode laser (OBIS-FP-647LX, Coherent, USA) combined with a two-axis AOD scanner (DTS-XY400-647, AA optoelectronics, France). The laser beam was directed to the back focal plane of a high-NA oil immersion objective (100× UPlanSApo, 1.4 NA, Olympus, Japan) using a pair of telecentric lenses (T1 and T2, *f* = 500 mm, AC254-500-A, Thorlabs, USA). The objective collected both reflected and scattered light fields, which were relayed to an sCMOS camera (PCO.edge 4.2, PCO, Germany) through a tube lens (TL1, *f* = 750 mm, AC254-750-A, Thorlabs, USA), the optical configuration of which resulted in a field of view of approximately 20 × 20 μm^2^.

For fluorescence imaging of GFP-labeled ER, APGs, and microtubules, an LED light source (SOLIS-3C, Thorlabs, USA) was used for excitation after the LED light was filtered with a bandpass filter (F1, FF02-482/18-25, Semrock, USA). A dichroic mirror (DM1: FF495-Di03, Semrock,) was used to combine the LED excitation into the main beam path to the sample dish. Fluorescence emission was guided by two dichroic mirrors (DM1 via transmission and DM2, FF552-Di02, Semrock, via reflection) and filtered through an emission bandpass filter (F2, FF02-520/28-25, Semrock, USA). The signal was then projected onto an EMCCD camera via a separate tube lens (TL2, *f* = 400 mm, AC254–400-A, Thorlabs, USA), expanding the field of view to approximately 24 × 24 μm^2^.

### iSCAT Image Processing

Details of the iSCAT image processing methods have been described previously.^22^ Briefly, iSCAT microscopy captures the interference between the scattering field (*E*_*s*_) from intracellular targets and the reference field (*E*_*r*_) reflected at the glass-medium interface. The detected intensity (*I*_det_) is expressed as

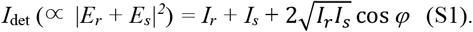

Here, *φ* is the relative phase between the fields. For small scatterers, the third interferometric term becomes more significant than the second pure scattering term, allowing the iSCAT contrast (*C*) to be approximately given as

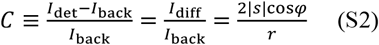

where *I*_back_ and *I*_diff_ are the background intensity and its difference from *I*_det_, and *r* and *s* are the reflection and scattering coefficients, respectively.

To obtain SBR-iSCAT images, a temporal-median cell image was first generated by averaging 20 (or 10) consecutive raw images acquired at 50 (or 200) Hz, and then a temporal-median background image was generated by averaging 3,000 (or 9,000) consecutive images taken from a cell-free zone. The differential image was obtained by subtracting the latter from the former. The resulting differential image was then divided by the background image (Eq. (S2)). On the other hand, a TD-iSCAT image was generated by dividing a temporal-median cell image – created from 20 (or 50) consecutive frames at 50 (or 200) Hz – by another such image taken after a 0.2 (or 0.125) s interval. iSCAT variance maps were generated by calculating pixel-wise standard deviations of TD iSCAT contrast for 200 (4 s) or 400 (2 s) consecutive TD-iSCAT images acquired at 50 and 200 Hz, respectively.

#### Measurement of The Instantaneous Speed and MSD of APG from APG Trajectories

Instantaneous speeds (*v*) were calculated from manually tracked trajectories in SBR-iSCAT image sequences acquired at 50 and 200 Hz (Figures 1A and 2F) using

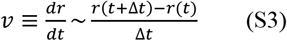

where *r*(*t*) is the APG position at time *t*, and *Δt* is given by *1/f* (Hz). To reduce fluctuations in *v* caused by thermal noise and pixel resolution limits (∼ 40 nm/pixel), *v* was calculated from 50-point (1 s) or 100-point (0.5 s) time-averaged trajectories from SBR-iSCAT images acquired at 50 or 200 Hz, respectively.

The MSD of APGs over a time interval *δt* is defined as the average squared displacement over *N* successive time lags:

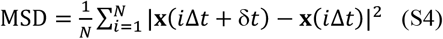

where **x**(τ) is the APG position at time *τ*.

### Statistical Analysis

The data presented in this work are representative of our experimental results. We performed iSCAT time-lapse imaging on four different COS-7 cells, including wild-type cells and those expressing GFP-labeled ER and GFP-labeled LC3. To validate the capability of iSCAT for label-free visualization of ER dynamics, we examined two conditions: wild type (Figure S2) and GFP-ER-expressing cells (Figure 1). For APG dynamics, two COS-7 cells expressing GFP-labeled LC3 were analyzed. In one of these cells, we observed the transformation of an APG from an elongated form into a closed sphere (Figure S3). A total of 22 APG trajectories were analyzed from the same field of view (20 × 20 μm^2^) within a single cell (Figure 2E). These trajectories were manually tracked from fluorescence image sequences acquired at 10 Hz. Of these, two representative APGs were further analyzed by manually tracking them in complementary SBR-iSCAT image sequences acquired at 200 Hz (Figures 2F and 4B). All figures were prepared using OriginPro (Origin lab), MATLAB (MathWorks), or Illustrator (Adobe).

## ASSOCIATED CONTENT

The Supporting Information is available free of charge on the ACS Publications website. Experimental setup of iSCAT integrated with fluorescence microscopy (Figure S1), iSCAT image processing methods for visualizing dynamic cytoplasmic components (Figure S2), visualization of APG transformation and ER remodeling events driven by APG transport (Figure S3), trajectories of 21 APGs interacting with the ER-microtubule networks during their journey (Figure S4), zoomed-in view of an APG trajectory, color-coded by time (Figure S5) and sequence of iSCAT variance maps illustrating the dynamic interplay between the APG and the surrounding ER network (Figure S6); legends for Movies S1-S11 (PDF).

Web Enhanced Objects: Movies

Label-free visualization of ER-cargo interaction via iSCAT microscopy (Movie S1), head-on collision of a dynamic cargo with a transverse ER tubule (Movie S2), ER tubule displacement by a dynamic cargo upon collision (Movie S3), ER hitchhiking on a dynamic cargo, leading to the formation of a new ER junction (Movie S4), transformation of an elongated LC3 structure into a spherical body (Movie S5), simultaneous visualization of APG dynamics, ER, and microtubule networks using single-color LC3 labeling (Movie S6), APG motion shift between random walk and directional drift near the ER-microtubule intersection (Movie S7), rapid short-range APG translocation at the ER-microtubule intersection after a slowly drifting phase (Movie S8), fast, long-range, directional transport of the APG along a microtubule (Movie S9), high-speed capture of the moment of APG transfer from the ER to an adjacent microtubule (Movie S10) and dynamic interplay between the APG and the surrounding ER network (Movie S11) (AVI).

## AUTHOR INFORMATION

### Author Contributions

Conceptualization: J.P., S.H., and M.C. Methodology: J.P., I.L., H.M., H.J. Investigation: J.P., S.H., and M.C., helped by I.L., H.M., and H.J.; M.L. and C.K. prepared GFP-LC3-expressed COS-7 cell lines. Funding acquisition: S.H. and M.C. Supervision: S.H. and M.C. Writing: J.P., S.H., and M.C. All of the authors received and approved the manuscript and Supporting Information.

### Notes

The authors declare no competing financial interest.

## ACKNOWLEDGMENTS

This work was supported by IBS-R023-D1 (MC), RS-2024-00347898 (SH), and RS-2023-00221182 (SH) through the NRF funded by the Ministry of Science and ICT.

## Notes

### Competing Interest Statement

The authors have declared no competing interest.

## REFERENCES

(1) Palade, G. Intracellular aspects of the process of protein synthesis. Science 1975, 189, 347–358.

(2) Schwarz, D. S.; Blower, M. D. The endoplasmic reticulum: structure, function and response to cellular signaling. Cell. Mol. Life. Sci. 2016, 73, 79–94.

(3) McNiven, M. A.; Thompson, H. M. Vesicle formation at the plasma membrane and trans-Golgi network: The same but different. Science 2006, 313, 1591–1594.

(4) Joshi, A. S.; Zhang, H.; Prinz, W. A. Organelle biogenesis in the endoplasmic reticulum. Nat. Cell Biol. 2017, 19, 876–882.

(5) Wenzel, E. M.; Elfmark, L. A.; Stenmark, H.; Railborg, C. ER as master regulator of membrane trafficking and organelle function. J. Cell Biol. 2022, 221, e202205135.

(6) Waterman-Storer, C. M.; Salmon, E. D. Endoplasmic reticulum membrane tubules are distributed by microtubules in living cells using three distinct mechanisms. Curr. Biol. 1998, 8, 798–806.

(7) English, A. R.; Zurek, N.; Voeltz, G. K. Peripheral ER structure and function. Curr. Opin. Cell Biol. 2009, 21, 596–602.

(8) Neefjes, J.; Jongsma, M. M. L.; Berlin, I. Stop or go? Endosome positioning in the establishment of compartment architecture, dynamics, and function. Trends Cell Biol. 2017, 27, 580–594.

(9) Guo, Y.; Li, D.; Zhang, S.; Yang, Y.; Liu, J.-J.; Wang, X.; Liu, C.; Milkie, D. E.; Moore, R. P.; Tulu, U. S.; Kiehart, D. P.; Hu, J.; Lippincott-Schwartz, J.; E. Betzig, Visualizing intracellular organelle and cytoskeletal interactions at nanoscale resolution on millisecond timescales. Cell 2018, 175, 1430–1442.

(10) Koppers, M.; Özkan, N.; Farias, G. G. Complex interactions between membrane-bound organelles, biomolecular condensates, and the cytoskeleton. Front. Cell Dev. Biol. 2020, 8, 618733.

(11) Qin, J.; Guo, Y.; Xue, B.; Shi, P.; Chen, Y.; Su, Q. P.; Hao, H.; Zhao, S.; Wu, C.; Yu, L.; Li, D.; Sun, Y. ER-mitochondria contacts promote mtDNA nucleoids active transportation via mitochondrial dynamic tabulation. Nat. Commun. 2020, 11, 4471.

(12) Lu, M.; van Tartwijk, F. W.; Lin, J. Q.; Nijenhuis, W.; Parutto, P.; Fantham, M.; Christensen, C. N.; Avezov, E.; Holt, C. E.; Tunnacliffe, A.; Holcman, D.; Kapitein, L.; Schierle, G. S. K.; Kaminski, C. F. The structure and global distribution of the endoplasmic reticulum network are actively regulated by lysosomes. Sci. Adv. 2020, 6, eabc7209.

(13) Gubas, A.; Dikic, I. ER remodeling via ER-phagy. Mol. Cell 2022, 82, 1492–1500.

(14) Valm, A M.; Cohen, S.; Legant, W. R.; Melunis, J.; Hershberg, U.; Wait, E.; Cohen, A. R.; Davidson, M. W.; Betzig, E.; Lippincott-Schwartz, J. Applying systems-level spectral imaging and analysis to reveal the organelle interactome. Nature 2017, 546, 162–167.

(15) Rose, G. G.; Pomerat, C. M. Phase contrast observations of the endoplasmic reticulum in living tissue cultures. J. Biophys. Biochem. Cytol. 1960, 8, 423–430.

(16) Buckley, I. K. Phase contrast observations on the endoplasmic reticulum of living cells in culture. Protoplasma 1965, 59, 569–588.

(17) Ma, Y.; Guo, S.; Pan. Y.; Pan, R.; Smith, Z. J.; Lane, S.; Chu, K. Quantitative phase microscopy with enhanced contrast and improved resolution through ultra-oblique illumination (UO-QPM). J. Biophoton. 2019, 12, e201900011.

(18) Lindfors, K.; Kalkbrenner, T.; Stoller, P.; Sandoghdar, V. Detection and spectroscopy of gold nanoparticles using supercontinuum white light confocal microscopy. Phys. Rev. Lett. 2004, 93, 037401.

(19) Kukura, P.; Ewers, H.; Müller, C.; Renn, A.; Helenius, A.; Sandoghdar, V. High-speed nanoscopic tracking of the position and orientation of a single virus. Nat. Methods 2009, 6, 923–929.

(20) Young, G.; Hundt, N.; Cole, D.; Fineberg, A.; Andrecka, J.; Tyler, A.; Olerinyova, A.; Ansari, A.; Marklund, E. G.; Collier, M. P. et al. Quantitative mass imaging of single biological macromolecules. Science 2018, 360, 423–427.

(21) Lee, I.-B.; Moon, H.-M.; Park, J.-S.; Lee, S.-H.; Lee, J.; Park, S. H.; Lee, S; Hong, S.-C.; Cho, M. Determination of the absolute concentration of Rayleigh particles via interferometric scattering microscopy. ACS Photonics 2025, 12, 3763–3771.

(22) Park, J.-S.; Lee, I.-B.; Moon, H.-M.; Joo, J H; Kim, K.-H.; Hong, S.-C.; Cho, M. Label-free and live cell imaging by interferometric scattering microscopy. Chem. Sci. 2018, 9, 2690−2697.

(23) Taylor, R. W.; Mahmoodabadi, R. G.; Rauschenberger, V.; Giessl, A.; Schambony, A.; Sandoghdar, V. Interferometric scattering microscopy reveals microsecond nanoscopic protein motion on a live cell membrane. Nat. Photonics 2019, 13, 480–487.

(24) Park, J.-S.; Lee, I.-B; Moon, H-M.; Ryu, J.-S.; Kong, S.-Y.; Hong, S.-C.; Cho, M. Fluorescence-combined interferometric scattering imaging reveals nanoscale dynamic events of single nascent adhesions in living cells. J. Phys. Chem. Lett. 2020, 11, 10233–10241.

(25) Hsiao, Y.-T.; Tsai, C.-N.; Chen, T.-H.; Hsieh, C.-L. Label-free dynamic imaging of chromatin in live cell nuclei by high-speed scattering-based interference microscopy. ACS Nano 2022, 16, 2774–2788.

(26) Park, J.-S.; Lee, I.-B.; Moon, H.-M.; Hong, S.-C.; Cho, M. Long-term cargo tracking reveals intricate trafficking through active cytoskeletal networks in the crowded cellular environment. Nat. Commun. 2023, 14, 7160.

(27) Park, J.-S.; Lee, I.-B.; Hong, S.-C.; Cho, M. Label-free interference imaging of intracellular trafficking. Acc. Chem. Res. 2024, 57, 1565–1576.

(28) Küppers, M.; Albrecht, D.; Kashkanova, A. D.; Lühr, J.; Sandoghdar, V. Confocal interferometric scattering microscopy reveals 3D nonoscopic structure and dynamics in live cells, Nat. Commun. 2023, 14, 1962.

(29) Mazaheri, M.; Kasaian, K.; Albrecht, D.; Renger, J.; Utikal, T.; Holler, C.; Sandoghdar, V. iSCAT microscopy and particle tracking with tailored spatial coherence. Optica 2024, 11, 1030–1038.

(30) Wu, B.-K.; Tsai, S.-F.; Hsieh, C.-L. Simplified interferometric scattering microscopy using low-coherence light for enhanced nanoparticle and cellular imaging. J. Phys. Chem. C 2025, 129, 5075–5085.

(31) Salogiannis, J.; Egan, M. J.; Reck-Peterson, S. L. Peroxisomes move by hitchhiking on early endosomes using the novel linker protein PxdA. J. Cell Biol. 2016, 212, 289–296.

(32) Salogiannis, J.; Reck-Peterson, S. L. Hitchhiking: A non-canonical mode of microtubule-based transport. Trends Cell Biol. 2017, 27, 141–150.

(33) Mogre, S. S.; Christensen, J. R.; Niman, C. S.; Reck-Peterson, S. L.; Koslover, E. F. Hitching a ride: Mechanics of transport initiation through linker-mediate hitchhiking. Biophys. J. 2020, 118, 1357–1369.

(34) Spits, M.; Heesterbeek, I. T.; Voortman, L. M.; Akkermans, J. J.; Wijdeven, R. H.; Cabukusta, B.; Neefjes, J. Mobile late endosomes modulate peripheral endoplasmic reticulum network architecture. EMBO Rep. 2021, 22, e50815.

(35) Yoon, M. J.; Choi, B.; Kim, E. J.; Ohk, J.; Yang, C.; Choi, Y. G.; Lee, J.; Kang, C.; Song, H. K.; Kim, Y. K. et al. UXT chaperone prevents proteotoxicity by acting as an autophagy adaptor for p62-dependent aggrephagy. Nat. Commun. 2021, 12, 1955.

(36) Tooze, S. A.; Yoshimori, T. The origin of the autophagosomal membrane. Nature 2010, 12, 831-835 (2010).

(37) Faller, E. M.; Villeneuve, T. S.; Brown, D. L. MAP1a associated light chain 3 increases microtubule stability by suppressing microtubule dynamics. Mol. Cell. Neurosci. 2009, 41, 85–93.

(38) Ross, J. L.; Ali, M. Y.; Warshaw, D M. Cargo transport: molecular motors navigate a complex cytoskeleton. Current. Opin. Cell Biol. 2008, 20, 41–47.

(39) Bálint, S.; Vilanova, I. V.; Álvarez, Á. S.; Lakadamyali, M. Correlative live-cell and super-resolution microscopy reveals cargo transport dynamics at microtubule intersections. Proc. Natl. Acad. Sci. U. S. A. 2013, 110, 3375–3380.

(40) Westrate, L. M.; Lee, J. E.; Prinz, W. A.; Voeltz, G. K. From follows function: The importance of endoplasmic reticulum shape. Annu. Rev. Biochem. 2015, 84, 791–811.

(41) Gurel, P. S.; Hatch, A. L.; Higgs, H. N. Connecting the cytoskeleton to the endoplasmic reticulum and Golgi. Curr. Biol. 2014, 24, R660–672.

